# Sub-second Extracellular Impedance Measurement of Epithelial Cell Monolayers using Step Excitations and Time-domain Analysis

**DOI:** 10.64898/2026.02.17.706376

**Authors:** Rongming Guo, Athena J. Chien, Jake Hawks, Benjamin Magondu, Bo Yang, Xavier Acevedo, Adrienne L. Watson, Bob Lewis, Chris Hatcher, Craig R. Forest

## Abstract

Extracellular electrochemical impedance spectroscopy (EIS) is emerging as a powerful technique in *in vitro* epithelial research, offering quantitative insights into barrier integrity, morphology, and apical-basolateral polarity noninvasively through metrics such as transepithelial electrical resistance (TER/TEER), transepithelial capacitance (TEC), and membrane ratio (*α*), respectively. However, due to the broad range of frequencies probed, EIS typically requires tens of seconds per measurement, limiting its ability to capture more rapid biological phenomena. We present Time-domain Epithelial Impedance Measurement (TEIM), a method for sub-second extracellular impedance measurements of epithelial cell monolayers based on step (Heaviside function) current excitation and time-domain analysis of the voltage transients, without the need for Fourier transforms. We experimentally demonstrate TEIM measuring TER, TEC, *α*, and model-derived impedance spectrum at ∼0.3 s sampling rate, which represents a 100- fold improvement in time resolution compared with traditional EIS. The accuracy and precision of TEIM were benchmarked against EIS on both electrical circuits and epithelial cell monolayers of immortalized Human Bronchial Epithelial (16HBE) and Human Colorectal Adenocarcinoma (Caco-2) (n = 3 for each), and average percent errors for TER, TEC, and *α* ranged from 0.17-3.55%, 1.13-8.96%, and 0.59-26.35%, respectively. Application of TEIM to monitor Caco-2 responses to saponin, a quick-acting pore-forming detergent, revealed smoothly gated double-exponential transient TER and TEC dynamics that were too rapid to be adequately captured previously using EIS. Overall, TEIM enables electrophysiology studies of rapid changes in epithelial cell culture models and possibly more complex *in vitro* models, holding promise for future applications in areas such as disease modeling, therapeutic development, and beyond.

## 1 Introduction

Epithelial tissues play a crucial role in the human body by regulating the transport of nutrients and waste products and defending against pathogens. The ability to monitor their barrier function and transport dynamics, particularly in real time, is therefore essential for applications such as disease modeling, early infection diagnosis, and therapeutics optimization.

Biological events that modulate epithelial barrier functions unfold over a wide range of durations, as shown in Figure 1 (A). Slow processes such as cell division and maturation [1, 2], as well as structural tissue damage due to chronic inflammation [2, 3] occur gradually over 10^3^ to 10^6^ seconds (hours to days); faster events such as neutrophil transepithelial migration [4, 5] and the functional remodeling of the tight junction (TJ) [6,7] alter barrier structure and function over 10^0^ to 10^3^ seconds (minutes to seconds); virtually instantaneous events such as cooperative ion channel gating [8, 9] can play out within sub-second intervals.

**Figure 1:**
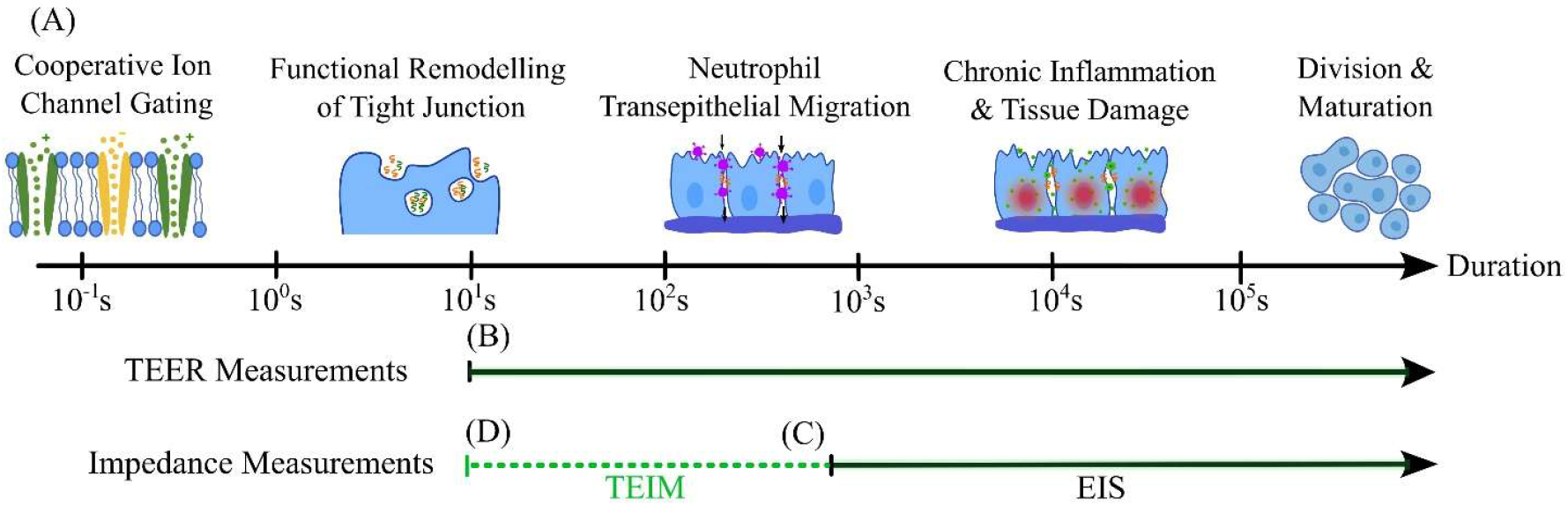
Duration (i.e., time constant) of biological events in epithelial cells (A) and time resolution of electrophysiology sensing technologies (B-D). TEER can be measured approximately every second [10], and is suitable for events unfolding on the same order (B). EIS-based impedance measurements, while providing richer biological information through both TEER/TER and TEC, can take over one minute [11, 12], limiting their applicability to slower events unfolding over minutes to hours (C). TEIM extends impedance measurement time resolution to the sub-second regime, allowing more rapid biological phenomena to be studied while preserving the information richness of EIS (D).

Various experimental methods have been used to study epithelial tissues *in vitro*, including optical techniques such as fluorescence microscopy, micro-optical coherence tomography (*μ*OCT), and Terahertz Attenuated Total Reflection (THz-ATR) [4, 13, 14]; chemical techniques such as tracer flux assays [15, 16]; and electrophysiological techniques such as short-circuited currents (I_sc_) measurements, transepithelial electrical resistance (TER/TEER) measurements [2, 3, 17–19], and electrochemical impedance spectroscopy (EIS) [11,12,20–23]. Compared to I_sc_ and TEER, in which direct current (DC) electrical stimuli are applied to the cells, EIS captures their responses to alternating current (AC) stimuli over a discrete set of frequencies ranging from several Hz to tens of kHz, and calculates a complex impedance spectrum, *Z*(*ω*) *ϵ* ℂ. Then, using equivalent circuit model fitting [11, 12, 22, 23], *Z*(*ω*) can be interpreted to reveal not only TER/TEER, but also transepithelial capacitance (TEC), a quantity with a promising link to membrane morphology, adhesion and dielectric composition [24–26], and membrane ratio (*α*), which reflects the permeability and structural asymmetries between the apical and basolateral surfaces of many epithelial tissues [11].

One major disadvantage of EIS, however, lies in its limited measurement speed. Depending on the number of frequencies and the number of periods repeated at each frequency, acquisition time of a single EIS impedance spectrum can range from tens of seconds to over a minute on customized biosensing chips or commercially available platforms such as the cellZscope^®^ (nanoAnalytics, Münster, Germany) and Autolab^®^ (Metrohm, Herisau, Switzerland) [12, 23, 27] – these are nearly an order of magnitude slower than TEER measurements, which take only a couple of seconds on instruments such as the EVOM^®^ Manual (World Precision Instruments (WPI), Sarasota, FL, USA) and ECIS^®^ (Applied Biophysics, Troy, NY, USA) [10]. As shown in Figure 1 (B) and (C), a tradeoff currently exists between the information richness of EIS-based impedance and the measurement speed of TEER.

Dating back to the 1870s, transient analysis in the time domain has been widely applied across engineering disciplines and life sciences to identify properties of unknown dynamical systems. Some examples include the classical mass-spring-damper (MSD) system in mechanical engineering, the Windkessel model for cardiovascular function assessment in clinical biology [29], and whole-cell patch clamp electrophysiology in neuroscience [30]. Motivated by the same principles, we showed that transitioning from frequency-domain (EIS) to time-domain analysis can significantly accelerate impedance measurements of epithelial cell monolayers. Specifically, by applying a step current excitation across the cell layer and observing its transient voltage response, impedance can be calculated without using Fourier transforms. More importantly, due to their intrinsic biological properties, epithelial membranes exhibit time constants (*τ* ≈ TER × TEC) on the order of milliseconds [11, 12, 21], allowing the responses to settle very rapidly and, as a result, enabling impedance measurements to be performed at sub-second timescales in real time. While similar concepts were proposed as early as 1946 by Teorell et al [31] and again in 1982 by Suzuki et al [32], these earlier efforts were limited by conceptual abstraction (Teorell), the need for intracellular electrodes (Suzuki), and, most importantly, a lack of digital processing power necessary for real-time application.

In this article, we introduce the theoretical framework behind and present a prototype implementation of the time-domain, step-excitation-based rapid impedance measurement method for epithelial tissues, which we term Time-domain Epithelial Impedance Measurement (TEIM). The prototype implementation, referred to as the TEIM-sensor, was integrated with a commercial extracellular electrophysiology chamber and achieved an average measurement interval of approximately 0.3 s, extending the range of biological phenomena accessible to impedance-based studies by nearly two orders of magnitude (Figure 1 (D)). The accuracy and precision of the TEIM-sensor were evaluated on both electrical circuit mimics and biological samples of Human Colorectal Adenocarcinoma (Caco-2) and immortalized Human Bronchial Epithelial Cells (16HBE), with average percent errors for TER, TEC, and *α* ranging from 0.17-3.55%, 1.13-8.96%, and 0.59-26.35%, respectively. Finally, to demonstrate the advantage of higher sampling rate, TEIM-sensor was used to continuously monitor the rapid impedance dynamics of Caco-2 cells subjected to apical exposure to saponin, revealing smoothly gated double-exponential transient TER and TEC dynamics for the first time.

## 2 Materials and Methods

### 2.1 Theories and Modeling

#### 2.1.1 Electrical Circuit Models of Epithelia

Ions and charged particles are transported through the epithelial membranes via specialized channels or electrically separated by the insulating lipid bilayer (Figure 2 (A)). In epithelial electrophysiology, these mechanisms are modeled using equivalent electrical circuits of resistors and capacitors. While a high-fidelity model distinguishes between apical/basolateral, and paracellular/transcellular ion transport pathways (Figure 2 (B)), its mathematical redundancy necessitates both extracellular and intracellular measurements for unique parameter solution, which is a delicate process and can be time-consuming. As a practical alternative, we use the RCRC model shown in Figure 2 (C), which absorbs the paracellular resistance into the apical and basolateral membrane resistances, requires extracellular electrodes only, and has been applied to Retinal Pigment Epithelium (RPE), 16HBE, and Caco-2 [11, 12, 22].

**Figure 2:**
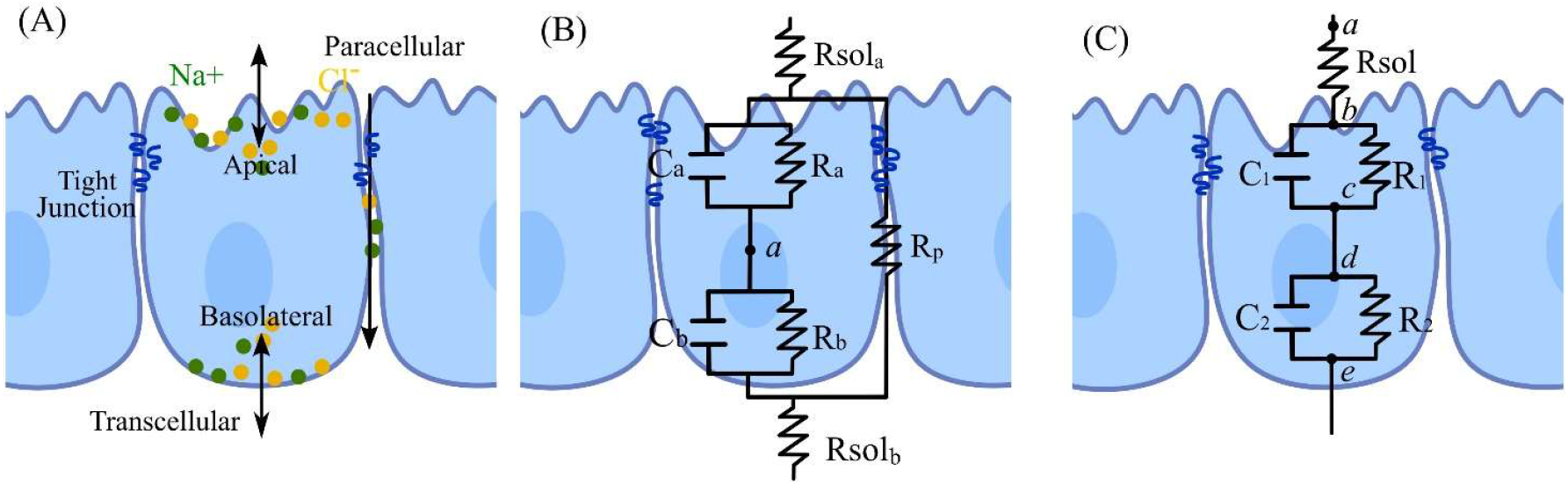
Physiological model (A) and electrical circuit models (B-C) of epithelia, modified from Chien et al. [11]. The physiological model describes behavior of ions and other charged particles, including conduction through transcellular and paracellular pathways and electrical separation by the lipid bilayer (A). The seven-parameter circuit model provides very high physiological fidelity but requires an intracellular measurement at point *a* to be uniquely resolved (B). The five-parameter (RCRC) model offers relatively high physiological fidelity while requiring only extracellular measurements at point *a* and *e* and has been applied to model RPE, 16HBE, and Caco-2 cells [11, 12, 20] (C).

Within the RCRC model, we define the bulk tissue impedance properties, including TER, TEC, and *α*, as follows:

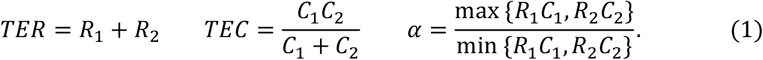

#### 2.1.2 Transient Transepithelial Voltage Response to a Step Current Excitation

Given that a step current stimulus is applied at time *t* = 0 across the epithelial cell layer using a pair of extracellular electrodes located at nodes a and e shown in Figure 2 (C), such that

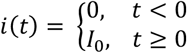

The resulting transepithelial voltage, measured by the same electrode pair, is denoted as *v*(*t*). To derive *v*(*t*), we write the governing ordinary differential equations (ODEs) for each circuit element and apply Kirchoff’s Current Law (KCL) at branching nodes *b* and *d*. First, across the solution resistance, the voltage follows Ohm’s Law,

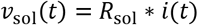

where *v*_sol_ is the voltage across node a and b. The current then splits at the apical junction (node *b*), either flowing through the apical membrane resistor *R*_1_ or charging the apical membrane capacitor *C*_1_:

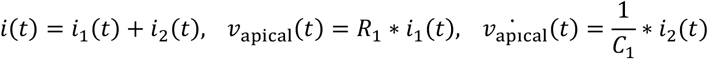

where *i*_1_ is the current passing through *R*_1_, *i*_2_ is the current charging *C*_1_, and *v*_apical_ is the voltage across node *b* and *c*. Similarly, for the basolateral membrane:

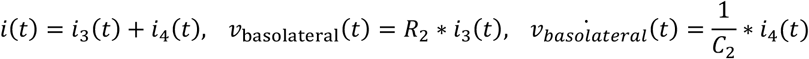

where *i*_3_ is the current passing through the basolateral membrane resistor *R*_2_, *i*_4_ is the current charging the basolateral membrane capacitor *C*_2_, and *v*_basolateral_ is the voltage across node *d* and *e*. Combined, the total transepithelial voltage is given by:

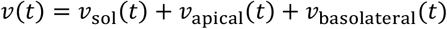

To analytically solve the above set of differential equations, we use Laplace transforms. First, we define the plant transfer function as

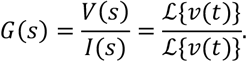

We then apply Laplace transforms to all the governing differential equations under zero initial conditions (enforced in Section 2.2.2). After rearranging the terms, the final transfer function is given by

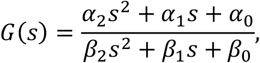

where

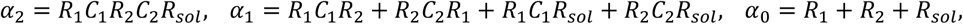

and

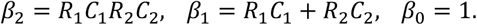

With *G*(*s*), the transient response of transepithelial voltage to a step current excitation can be calculated using the following inverse Laplace transform,

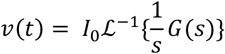

Solving analytically (Mathematica, Wolfram Research Inc., Champaign, IN, USA),

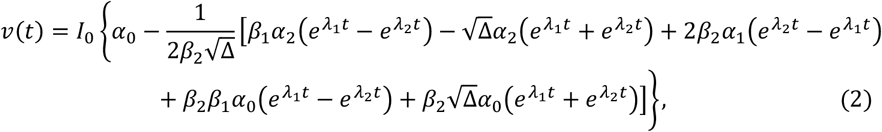

with two poles being

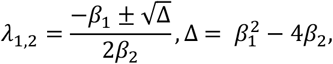

Which can be further simplified into

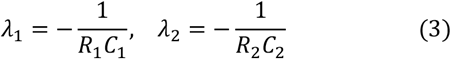

Since both poles are negative and real, the epithelial cell layer, as modelled here, is a stable, overdamped second-order system. Thus, the step response of the transepithelial voltage exhibits an asymptotically rising, non-oscillatory, double-exponential profile. A more detailed analysis of how changes in TER, TEC, and *α* affect the *v*(*t*) waveform is presented in Section 3.1.

#### 3.1.3 Estimating Circuit Parameters with Numerical Fitting

The circuit parameters of the RCRC model can be directly estimated from the measured transepithelial voltage step response using numerical curve fitting, without the need for Fourier transforms. First, we group the five unknown resistance and capacitance values into a parameter vector **p** such that

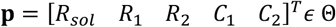

where Θ *∈* ℝ^5^ corresponds to the parameter space. To explicitly distinguish the measured transepithelial voltage response from its theoretical counterpart, and to account for discrete-time sampling in the actual implementation, we modify the notation of the measured response to

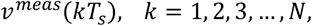

where *T*_*s*_ is the sampling period, *k* is the sample index, and *N* is the total number of samples. The corresponding theoretical response is denoted

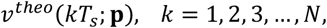

which follows the form given in Equation 2, is evaluated at the same sampling instants as the measured response, and is parameterized by **p**.

A numerical least-squares optimization algorithm then iteratively searches for the optimal set of circuit parameters **p**^∗^ that minimizes the sum of squared residuals (SSR) between the measured and modeled response such that

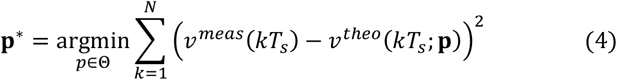

From **p**^∗^, the bulk tissue parameters are computed according to Equation 1.

#### 2.1.4 Model-derived Impedance Spectrum

For researchers already familiar with the Nyquist (Cole-Cole) plot representation of the impedance spectrum, such information can be derived from the fitted circuit parameters according to

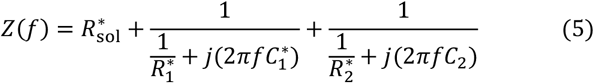

Which applies the series and parallel impedance combination rules to the RCRC circuit model. Here, *Z ϵ* ℂ is the impedance, *f ϵ* ℝ is the frequency point(s) of interest in Hz, and 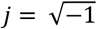 is the imaginary unit.

### 2.2 Instrumentation

#### 2.2.1 Hardware Architecture

We present a prototype implementation of TEIM, referred to as the TEIM-sensor. As shown in the functional schematic in Figure 3 (A), the TEIM-sensor consists of a National Instruments (NI) USB-6451 Data Acquisition (DAQ) device (National Instruments, Austin, Texas, USA), an HP development computer with an Intel Core Ultra 9 185H CPU, a USB 3.1 Gen 2 SuperSpeed+ Cable, and a custom galvanostatic driver (WPI). At its core, the galvanostatic driver integrates a voltage-controlled current source (VCCS) and two differential voltage-sense outputs, OSCI and OSCII. The current output of the VCCS is adjusted with respect to an on-board 10 *k*Ω reference resistor (*R*_ref_) such that

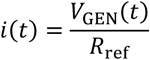

where *V*_GEN_ is the voltage control signal. In Figure 3 (B), an EndOhm (WPI) extracellular electrophysiology chamber with a transwell^®^ (Corning, Corning, NY, USA) insert of living cells serves as the device under test (DUT). VCCS and R_ref_ further connect in series with the DUT, delivering the regulated current through. Meanwhile, OSCI measures the combined voltage across both R_ref_ and DUT, and OSCII measures the voltage across R_ref_ alone, such that

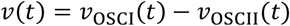

**Figure 3:**
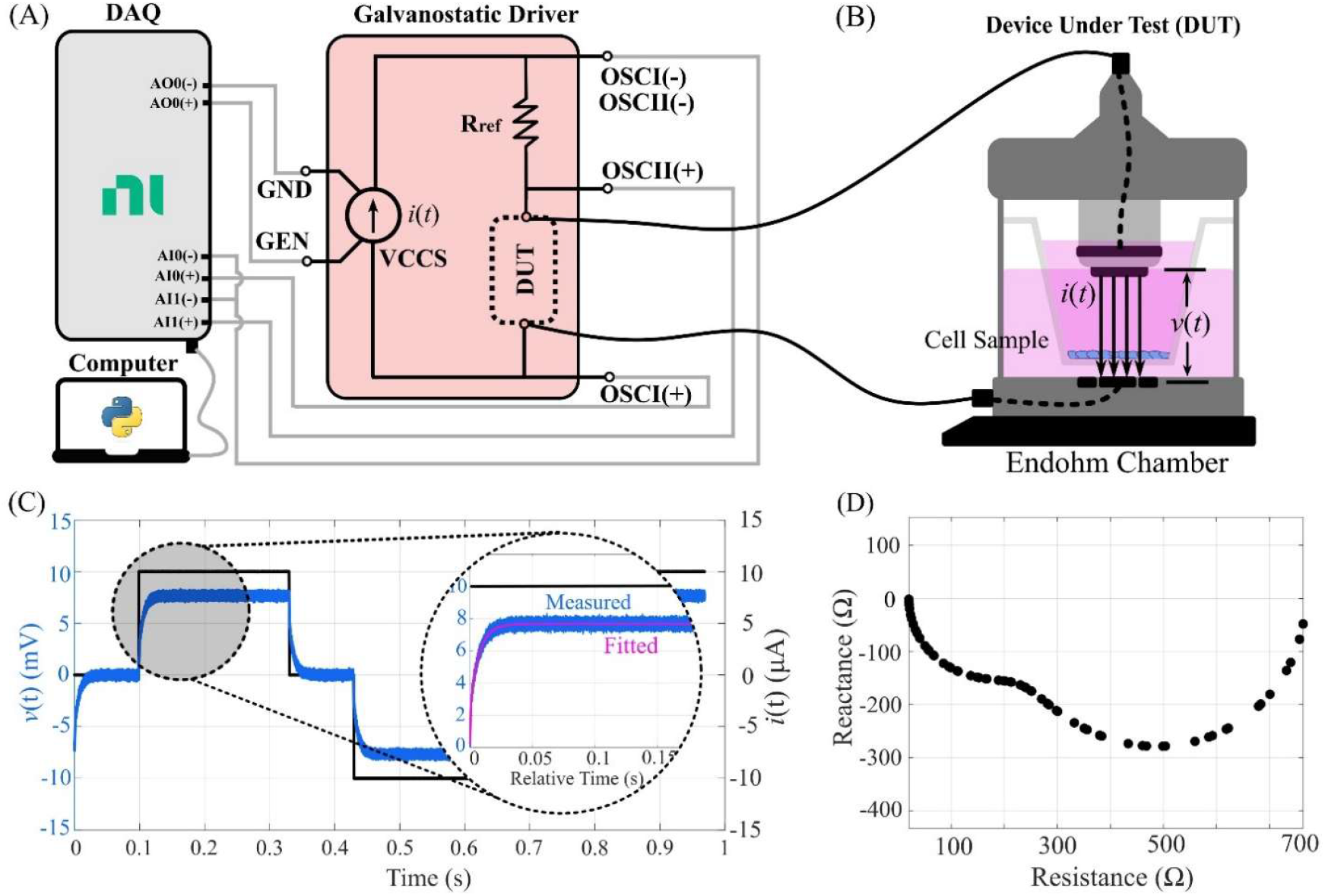
Hardware architecture (A-B) and measurement workflow (C-D) of the Time-domain Epithelial Impedance Measurement (TEIM)-sensor. Gray lines indicate essential internal wiring of the TEIM-sensor (A). An EndOhm electrophysiology chamber with a transwell of living epithelial cells serves as the device under test (DUT), and is connected to the galvanostatic driver circuit as indicated by the solid black lines (B). Step current with reversing polarities induces transient voltage responses of the cell layer, which is fitted to reveal the circuit parameters underneath (C). A Nyquist impedance spectrum can then be reconstructed using the fitted circuit parameters according to Equation 5 (D).

The control signal at GEN is provided by the DAQ through its analog output (AO) channel 0, and the voltage measurements at OSCI and OSCII are recorded by analog input (AI) channels 0 and 1. Communication between the DAQ and the development computer is carried out through a USB 3.1 SuperSpeed+ cable that supports the DAQ’s 1MS/s high sampling rate.

#### 2.2.2 Software and Configurations

A Python^®^ (version 3.13.2) program manages DAQ, data processing, and real-time data visualization. The step current magnitude *I*_0_ is set to 10*μ*A, standard for EIS or TEER implementations and safe for most epithelial cells [11, 12, 34]. The DAQ’s AO channel operates at its maximum update rate of 250 kS/s, while the AI channels operate at 98% of their maximum sampling rate at 980kS/s. An oversampling (block-averaging) ratio of 49:1 is applied to AI channels to reduce Gaussian white noise, resulting in a final effective sampling rate of 20 KS/s.

Before measuring, the TEIM-sensor needs to be calibrated using a blank transwell insert. VCCS first commands 0*μ*A, and the voltages at ‘OSCI’ (AI0) and ‘OSCII’ (AI1) are captured once the waveforms stabilize. These non-zero voltages, typically ranging from several mV to the low tens of mV, reflect the combined hardware offsets arising from the EndOhm chamber, galvanostatic driver, and the DAQ, and are compensated in subsequent measurements. As an optional step to improve the accuracy of R_sol_ estimation during curve fitting, VCCS can then output 10*μ*A to independently calculate of R_sol_ using Ohm’s law.

A representative measurement workflow is shown in Figure 3 (C). Each impedance measurement cycle consists of two consecutive phases: data acquisition (precisely 0.2 s) and analysis (roughly 0.1 s). During the first half of the data acquisition phase (e.g., from *t* = 0 s to *t* = 0.1 s), the VCCS first outputs *i*(*t*) = 0*μA* (black) to force all capacitive elements of the DUT to fully discharge; then, during the second half of the data acquisition phase (e.g., from *t* = 0.1 s to *t* = 0.2 s), the VCCS current is stepped to *i*(*t*) =10*μ*A, inducing a transient voltage step response *v*(*t*) (blue) across DUT.

The analysis phase begins afterwards. First, as shown in the zoomed-in window in Figure 3 (C), the measured response is input to a numerical fitter that iteratively solves the optimization problem defined in Equation (4) to estimate the underlying circuit parameters. TER, TEC, and *α* values are calculated according to Equation (1). Finally, as shown in Figure 3 (D), a Nyquist impedance spectrum is reconstructed from the solved circuit according to Equation (5) at frequency locations of interest before the next measurement cycle begins. Meanwhile, to minimize polarization of both the cell sample and the electrodes, the direction of the current is reversed between successive cycles.

### 2.3 Materials and Reagents

#### 2.3.1 Electrical Circuit Mimics

To isolate biological variations inherent to real epithelial cells and facilitate benchtop testing of the TEIM-sensor hardware, electrical circuit mimics of epithelial cell monolayers are constructed on a breadboard with axial-lead metal film resistors and aluminum electrolytic capacitors following the RCRC topology described in Section 2.1.1, Figure 2 (C). Component values are picked within the reported [11,12,21] and observed ranges of the cell type being modelled.

#### 2.3.2 Cell Culture

The epithelial cell lines used in this study include Caco-2 and 16HBE, both of which have been well studied with existing electrophysiology technologies such as TEER and EIS [2, 12]. Caco-2 cells were purchased from the American Type Culture Collection (ATCC HTB-37TM) frozen at passage 15 and cultured in T-25 flasks according to a protocol provided by SynVivo Inc. Huntsvile, AL, USA. The complete culture medium consisted of 79% RPMI 1640 medium (ATCC Modification, ThermoFisher Scientific, catalog no. A1049101), 20% fetal bovine serum (FBS) (ThermoFisher, catalog no. A3840002) and 1% Penicillin/Streptomycin (Pen/Strep, Gibco 15070-063). 16HBE cells were cultured following the protocol published by the CFFT lab [35] except that flasks were not coated with collagen. The complete culture medium was composed of 90% MEM (Gibco, catalog no. 11095-080), 10% FBS (Corning, catalog no. 35-015-CV), and the same 1% Pen/Strep. In addition, cells were cultured for at least 1 week prior to plating for experiments and used up to 12 weeks after thawing from frozen aliquots. Cells were plated on 1.12 cm^2^, 0.4 *μ*m pore size transwells^®^ (3460, Corning) at a density of roughly 250k cells (16HBE) and 180k cells (Caco-2) per well and maintained at liquid-liquid interface with 0.5 mL medium in the apical chamber and 1 mL in the basolateral chamber. Media changes were performed three times a week, and cells were passaged upon reaching >80% confluency.

#### 2.3.3 Saponin

Saponin is a naturally occurring amphipathic glycoside that can rapidly increase the permeability and alter cytoskeletal structure of epithelial cells by binding to their cholesterol-rich lipid bilayers and forming non-specific pores, leading to abrupt changes in both TER and TEC on epithelial cells [12, 19]. In this study, saponin is applied to Caco-2 cells to create a rapid cellular event to be measured by TEIM-sensor. To prepare the stock saponin solution, 0.05 g of saponin powder (ThermoFisher Scientific, catalog no. A18820.14) was dissolved in pre-warmed (37.5°C) complete Caco-2 culture medium to a final volume of 10 mL, yielding a concentration of 0.5% w/v, or 5 mg/mL. During experiments, the stock solution was further diluted upon addition to the apical chamber of the transwell.

## 3 Results and Discussion

### 3.1 Transepithelial Voltage Step Response

We demonstrate the effect of TER, TEC, and *α* on the time-domain transepithelial voltage step response, *v*(*t*), in Figure 4. The baseline circuit is designed to start with

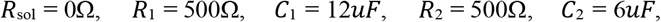

which corresponds to TER = 1000Ω, TEC = 4 *μ*F, and *α* = 2, values that fall within the normal range reported and observed for healthy epithelial monolayers. *I*_0_ is set to 10 *uA*, consistent with the TEIM-sensor implementation. The trends generalize to any current magnitude under which epithelial cell behavior remains well described by the circuit model.

**Figure 4:**
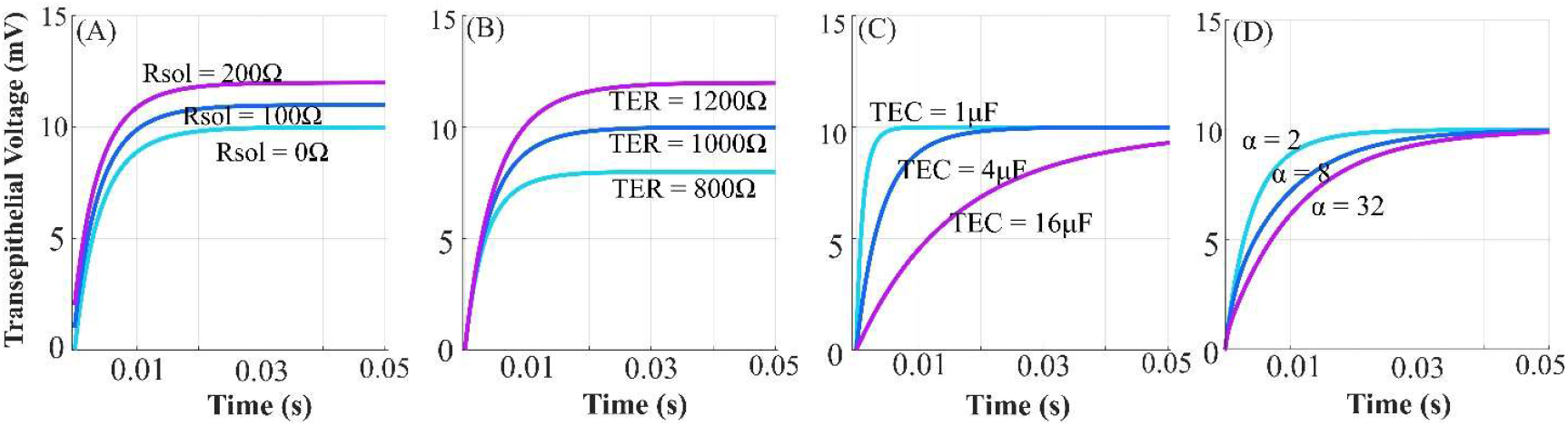
Effect of R_sol_, TER, TEC, and *α* on the transepithelial voltage step response waveform. Solution resistance corresponds to the y-intercept (A). TER, together with R_sol_, determines the steady state value (B). TEC governs the overall system response speed (C). *α* controls the degree to which the response exhibits second-order behavior (D).

As shown in subplot (A), the solution resistance R_sol_ is directly proportional to the y-axis intercept, or the initial value, of *v*(*t*). As shown together in subplots (A-B), the combined resistance of the solution and the cell determines the steady-state value. Indeed, evaluating Equation (2) at *t* = 0 and *t* → ∞ yields

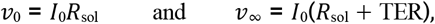

and the second equation reveals how TEER is traditionally calculated through “Ohm’s Law”.

The effects of TEC and *α* are less intuitive, as *v*(*t*) reflects the combined effect of two exponential modes. As shown in subplot (C), TEC is inversely related to the response speed of the system, with higher TEC increasing the time required to reach steady state. *α* can be interpreted as the relative speeds of the exponential modes. As shown in subplot (D), an *α* value closer to 1 suggests relatively balanced speeds, making the transient response appear nearly first-order; greater *α*, on the other hand, implies one mode is significantly faster than the other, giving the transient response a second-order appearance with a clearer change in slope early in the rise.

### 3.2 Accuracy and Precision

We evaluated the precision and accuracy of the TEIM-sensor on both electrical circuit mimics and biological samples of 16HBE and Caco-2 cells.

For electrical circuit mimics, the ground-truth TER, TEC, and *α* values were measured using a FLUKE 289 True-RMS Multimeter (Fluke Corporation, Everett, WA, USA), and the corresponding impedance spectra were reconstructed according to Equation (5). Following the ground-truth measurements, triplicate TEIM-sensor measurements were performed once every minute. Three replicate circuits were tested for each cell.

For biological samples, ground-truth values were obtained via the galvanostatic EIS protocol by Chien et al. [11] using a Metrohm Multi Autolab potentiostat/galvanostat electrochemical workstation (Metrohm AG, Herisau, Switzerland). Measured impedance spectra were fitted to the RCRC circuit model to extract TER, TEC, and *α*. Prior to measurements, cells were allowed to stabilize outside of the incubator for 15 minutes to mitigate transient impedance drift due to abrupt changes to the surrounding temperature and CO_2_ levels. Triplicate ground truth (EIS) and TEIM-sensor measurements were then performed in an alternating sequence for each sample (EIS 1^st^ → TEIM-sensor 1^st^ → EIS 2^nd^ → …). To match the ∼1-minute EIS duration, corresponding TEIM-sensor data were acquired continuously for 1 minute, yielding ∼180 data points.

Table 1 summarizes the accuracy (percentage error relative to the ground truth) and precision (coefficient of variation among triplicate measurements) of the TEIM-sensor. Overall, the TEIM-sensor performed better on electrical circuit mimics than biological samples, with 16HBE circuit mimics yielding percentage errors below 2% and coefficients of variation below 1% across TER, TEC, and *α*. TER percentage error remained below 5% for all groups, whereas TEC percentage error reached 8.96% for 16HBE biological samples. Furthermore, *α* accuracy and precision were significantly higher for 16HBE than for Caco-2 for both electrical circuit mimics and biological samples.

**Table 1:**
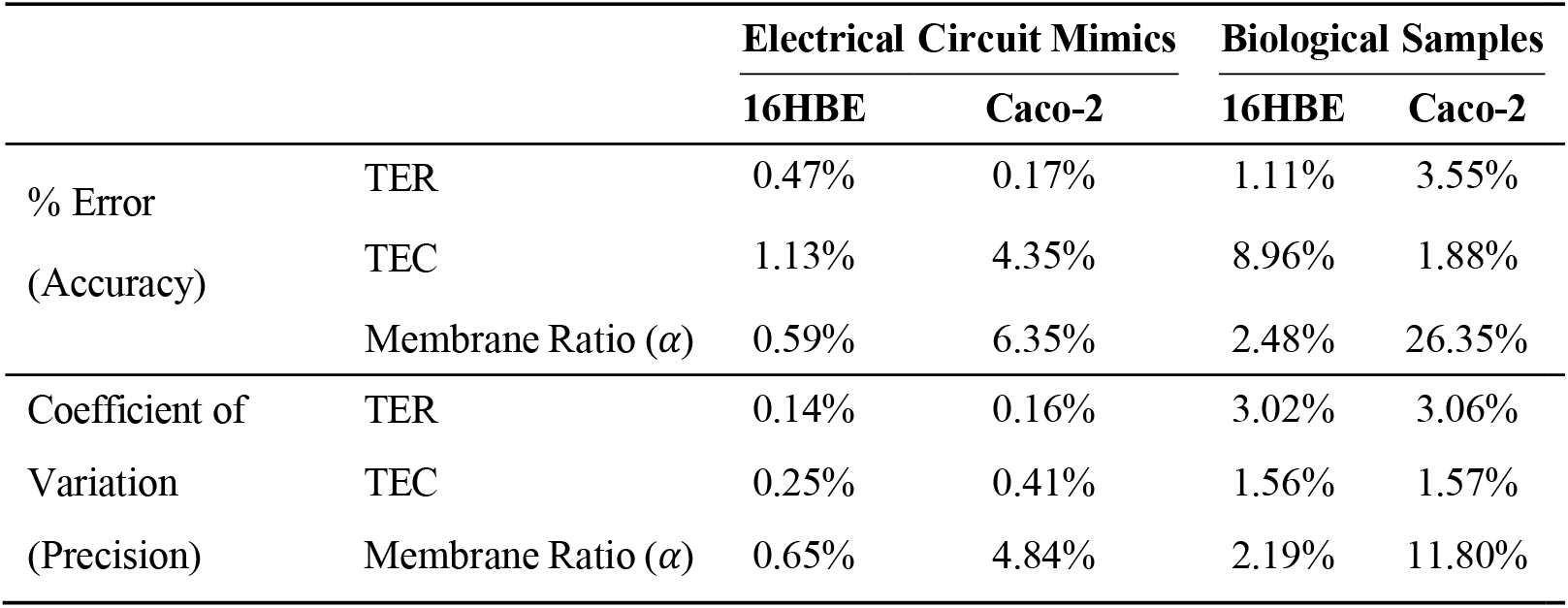
Accuracy and precision of the TEIM-sensor on both electrical circuit mimics and biological samples of 16HBE cells and Caco-2 cells. Accuracy is reported as the absolute percentage error between the TEIM-sensor measurements and the ground-truth values. Precision is reported as the coefficient of variation across triplicate TEIM-sensor measurements. Values were averaged across subject replicates within each subject group.

|  |  | Electrical Circuit Mimics |  | Biological Samples |  |
| --- | --- | --- | --- | --- | --- |
|  |  | 16HBE | Caco-2 | 16HBE | Caco-2 |
| % Error<br>(Accuracy) | TER | 0.47% | 0.17% | 1.11% | 3.55% |
|  | TEC | 1.13% | 4.35% | 8.96% | 1.88% |
| | Membrane Ratio ($\alpha$) | 0.59% | 6.35% | 2.48% | 26.35% |
| Coefficient of<br>Variation<br>(Precision) | TER | 0.14% | 0.16% | 3.02% | 3.06% |
|  | TEC | 0.25% | 0.41% | 1.56% | 1.57% |
| | Membrane Ratio ($\alpha$) | 0.65% | 4.84% | 2.19% | 11.80% |

Figure 5 provides absolute measurement details and a comparison between Nyquist impedance spectra for biological Caco-2 samples (plots for the remaining groups are available in the Supplemental Materials). As shown, despite the degradation of accuracy and precision of *α* for Caco-2, Nyquist impedance spectra (subplot (B)) show close agreement between the ground-truth and TEIM-sensor. Therefore, we hypothesize that this degradation arises from the limitations in numerical fitter, where the parameter space (Θ *∈* ℝ^5^) landscape becomes increasingly flat near the Caco-2 region due to its “single-bump” characteristic, making precise convergence more challenging.

**Figure 5:**
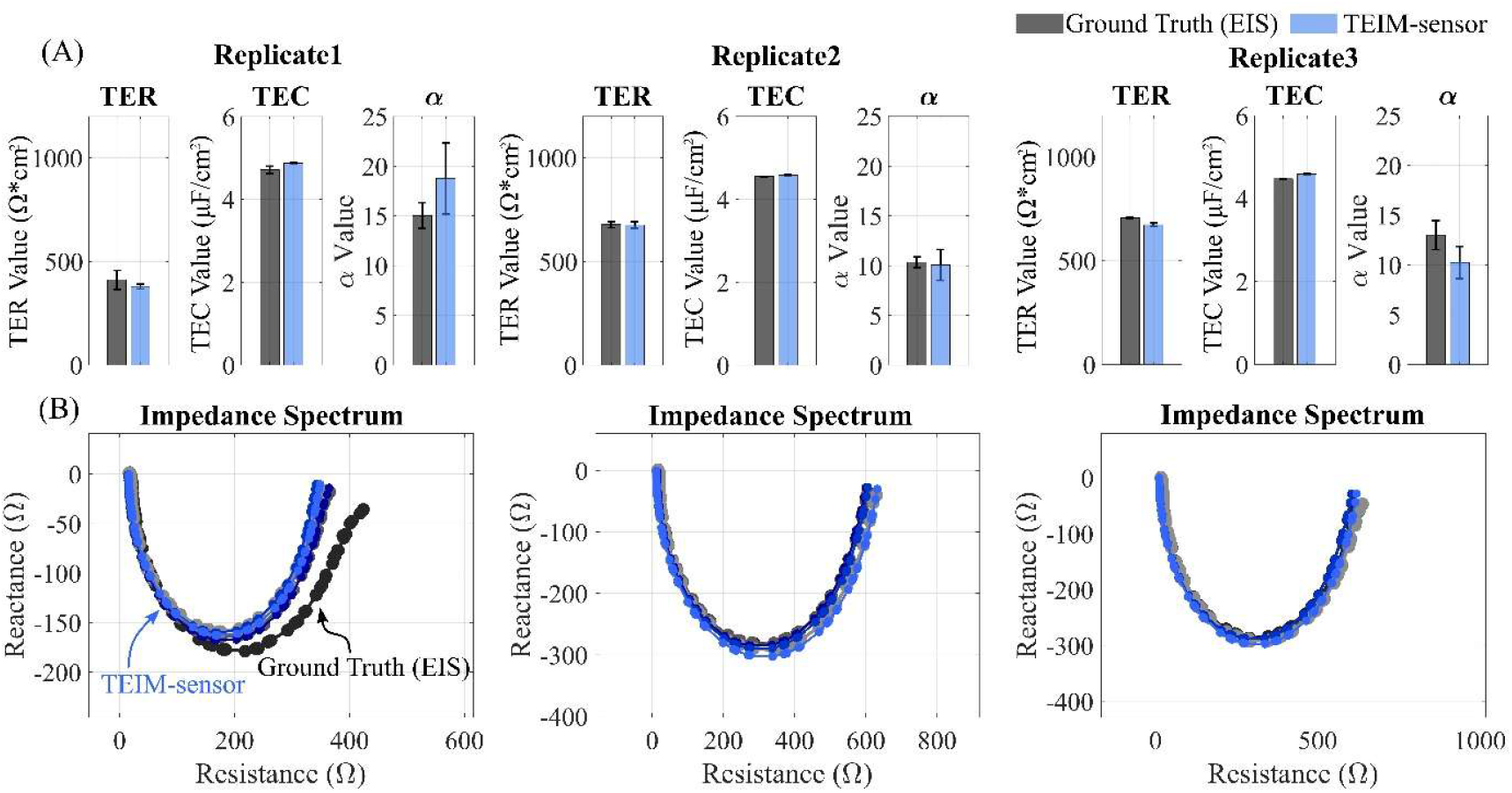
Accuracy and precision of TEIM-sensor on Caco-2 cells. Mean values (bars) and standard deviations (whiskers) of both ground truth EIS (gray) and TEIM-sensor (blue) measurements of TER, TEC, and *α* are shown in (A). Impedance spectra directly measured by ground truth EIS (gray) and reconstructed from TEIM-sensor measurements (blue) are overlaid in (B). TEIM-sensor Nyquist impedance spectra were reconstructed using parameters averaged over each 1-minute acquisition interval. Each column corresponds to one biological replicate, with triplicate measurements performed per replicate.

### 3.3 Continuous Monitoring of Caco-2 Exposure to Saponin

We highlight the advantages of the increased sampling rate enabled by the TEIM-sensor by monitoring the rapid TER and TEC dynamics of Caco-2 monolayers during apical exposure to saponin.

Six replicate samples were cultured to maturation (TER≈1200 Ω/cm2 and TEC≈4.5 *μ*F/cm2), with three assigned to the experimental group and the rest to the control. Upon removal from the incubator, cells were stabilized in the Endohm chamber for 5 minutes before 12 *μL* of 0.5% w/v stock saponin solution (experimental) or medium (control) were added to the apical chamber. This resulted in an effective final saponin concentration of 120*μ*g/mL in the experimental group, a value within the range reported in previous studies, including 10-40 *μ*g/mL at the lower end by Narai et al. [19] and 500 *μ*g/mL at the higher end by Linz et al. [12]. Immediately following addition, the TEIM-sensor continuously acquired impedance data for approximately 15 minutes.

Relative TER and TEC (defined as the measured values normalized to the first value recorded) time traces are shown in Figure 6 (A-B). Data were sampled every 0.31s on average and processed with a 10-sample moving-average filter (spanning roughly 3 s). Following saponin addition, relative TER in the experimental group declined sharply after a brief initial ramp-up, before reaching an inflection point after approximately 1-2 minutes and slowly approaching a final value close to 0.05. On the other hand, relative TEC in the experimental group exhibited longer ramp-up and increased asymptotically to a final value between 2-3. Based on the observations, we fitted the traces using double-exponential functions modified by a 3rd order polynomial soft gate:

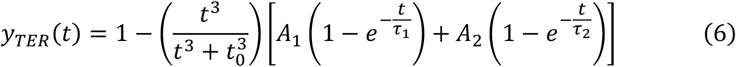

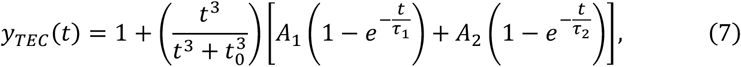

**Figure 6:**
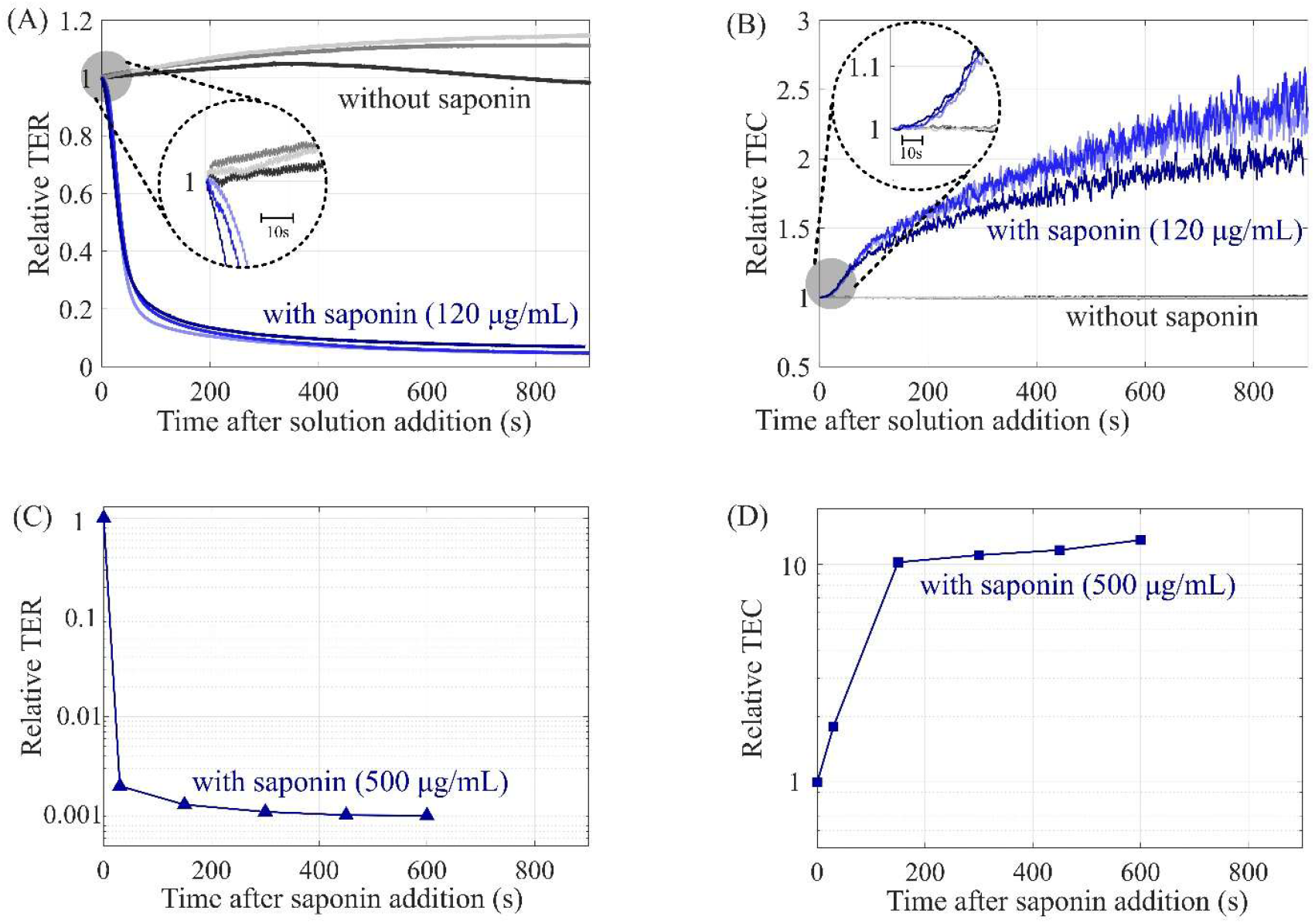
Time traces of relative TER and TEC dynamics of Caco-2 in response to apical exposure of Saponin. Subplots (A-B) show data associated with a saponin concentration of 120*μ*L/mg, measured using the TEIM-sensor with an average sampling interval of 0.3 s. Subplots (C-D) show data associated with a saponin concentration of 500*μ*L/mg, measured using EIS with an average sampling interval of 120 s; these data, including the use of logarithmic scales, are reproduced from [12] in their entirety.

where *t*_0_ reflects the duration of the initial onset, 1∓(*A*_1_ + *A*_2_) corresponds to the final value, and *τ*_1_ and *τ*_2_ represents the time constant of two exponential modes. Curve fitting was performed using MATLAB^®^ (Mathworks Inc. Natick, MA, USA) optimization toolbox. The resulting fitted expressions, averaged across samples, are:

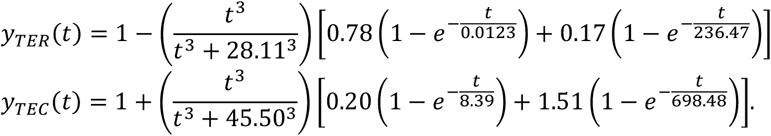

Several tentative biological interpretations can be drawn from these expressions. First, because the *t*_O_ (28.11s versus 45.50s), *τ*_1_ (0.0123s versus 8.39s), and *τ*_2_ (236.47s versus 698.48s) values differ significantly between TER and TEC dynamics, saponin likely influenced the permeability and morphology of the cell layer through two distinct pathways. In addition, the presence of two exponential modes with significantly different time constants for both TER and TEC dynamics suggests the possibility of further sub-processes within each pathway. While speculative, these interpretations demonstrate the possibilities for scientific discovery brought forth by TEIM.

For a sampling rate comparison, Figure 6 (C-D) presents data from Linz et al., who monitored the same process at a higher saponin concentration of 500*μ*g/mL using EIS and STX-2 electrodes [12]. Although similar exponential decay and asymptotic rise trends are visible, the lower sampling rate of EIS limited the level of dynamic detail that can be extracted. As a result, the multi-stage multi-mode curve-fitting analysis presented above might not have been possible, and the associated biological interpretations might never have been drawn.

Notably, our 120 *μ*g/mL concentration was optimized to balance biological response speed and signal-to-noise ratio; a higher concentration at 500*μ*g/mL caused an excessive drop of TER to single-digit ohms, rendering the transepithelial voltage signal indistinguishable from the electrical noise in the system, subsequently causing the fitter to fail.

## 4 Conclusions

This study presents an alternative method for extracellular impedance measurements of epithelial cell monolayers based on step current excitations and time-domain analysis of the transient voltage responses. Due to epithelial membranes’ inherently low resistances and capacitances, such transient responses settle extremely fast, allowing measurements to be performed at sub-second rates – nearly two orders of magnitude faster than the frequency-domain-based EIS. The accuracy and precision results are promising, and a representative real-world application shows its potential to facilitate scientific discoveries. In summary, this method enables continuous impedance monitoring of rapid cellular changes that are difficult to resolve using other systems, such as EIS, due to limited sampling rates, and establishes a foundation for future works in the space of epithelial electrophysiology.

### Declaration on the Use of Generative AI

During the preparation of the manuscript the author(s) used ChatGPT (openAI) and Gemini (Google) to polish the language quality and readability, primarily through grammar checks and word/phrase choice refinements. In addition, during the research of the background of the related technical fields, specifically epithelial electrophysiology, chatGPT with deep research were used to more efficiently synthesize information and organize previously published scholarly articles. After using this tool/service, the author(s) reviewed and edited the content as needed and take(s) full responsibility for the content of the published article.

## Supplemental Materials

**Figure S1:**
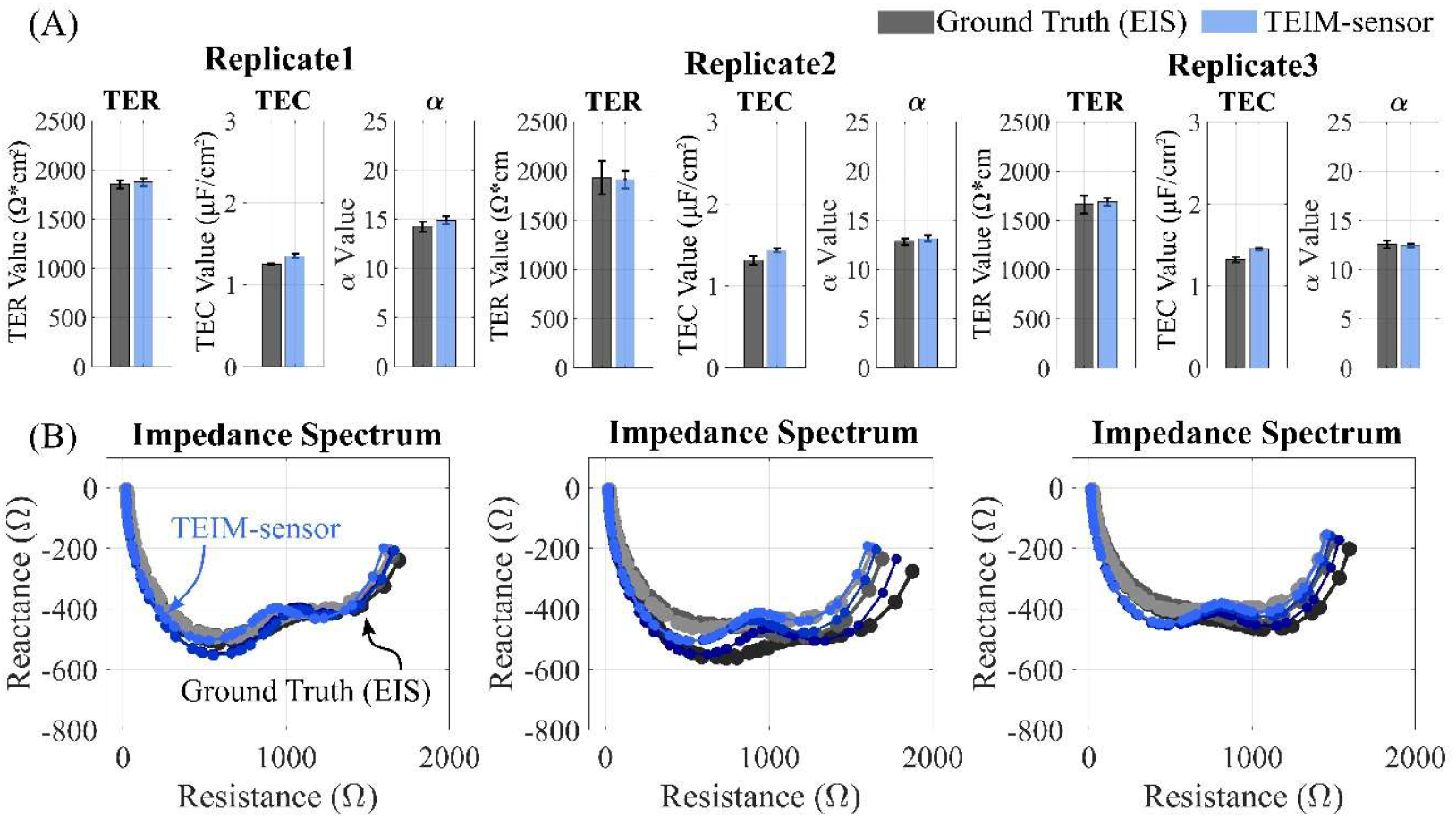
Accuracy and precision of TEIM-sensor on 16HBE cells. Data presentation and interpretation are the same as in Figure 5.

**Figure S2:**
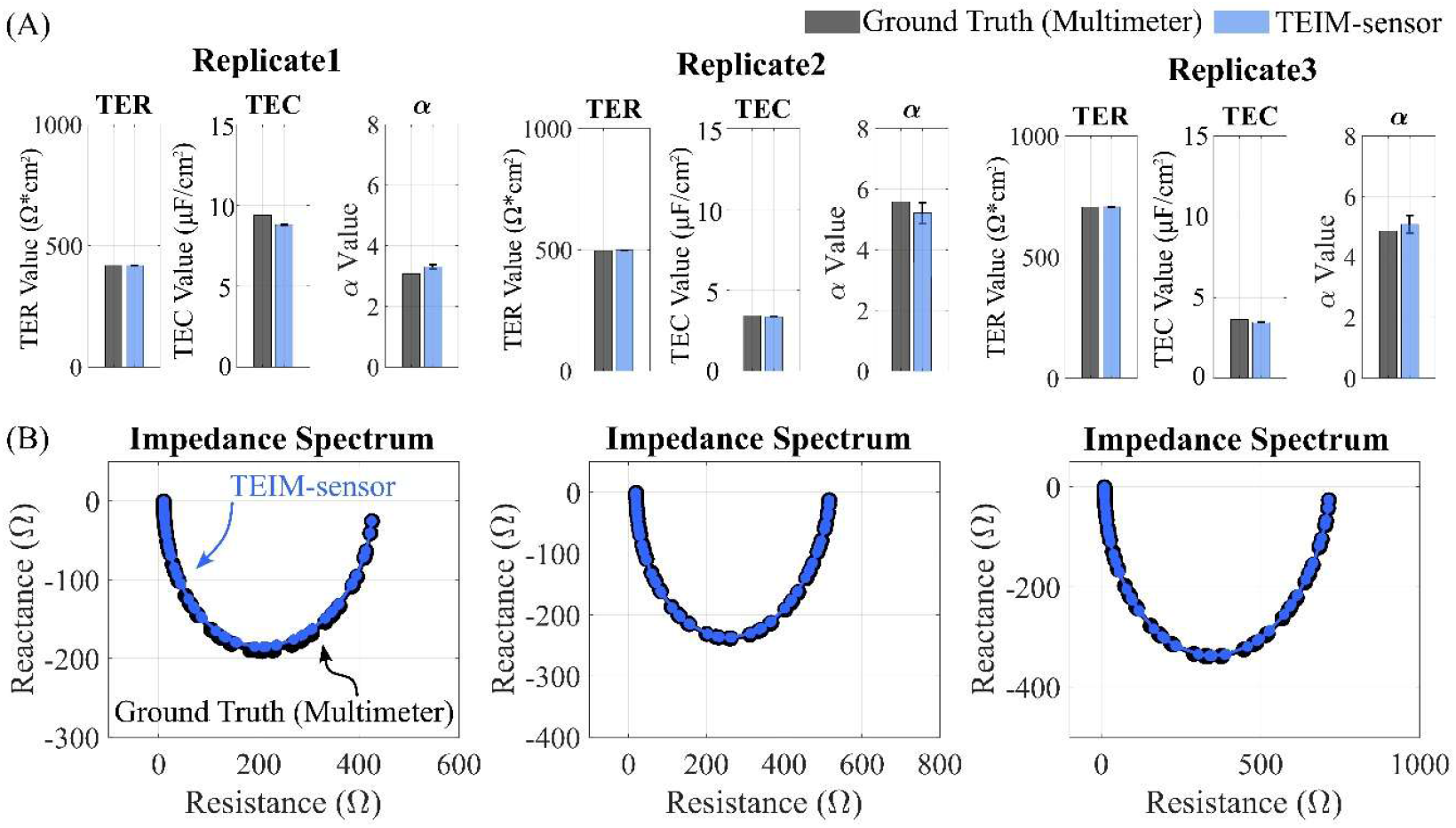
Accuracy and precision of TEIM-sensor on electrical circuit mimics of Caco-2. Ground truth measurements (gray) were performed once with a multimeter. Ground truth measurements (gray bars), as well as the mean values (blue bars) with standard deviations (whiskers) of TEIM-sensor measurements of TER, TEC, and *α* are shown in (A). Impedance spectra reconstructed from ground truth multimeter (gray) and TEIM-sensor (blue) measurements are overlaid in (B). Each column corresponds to one circuit replicate, with triplicate measurements performed per replicate.

**Figure S3:**
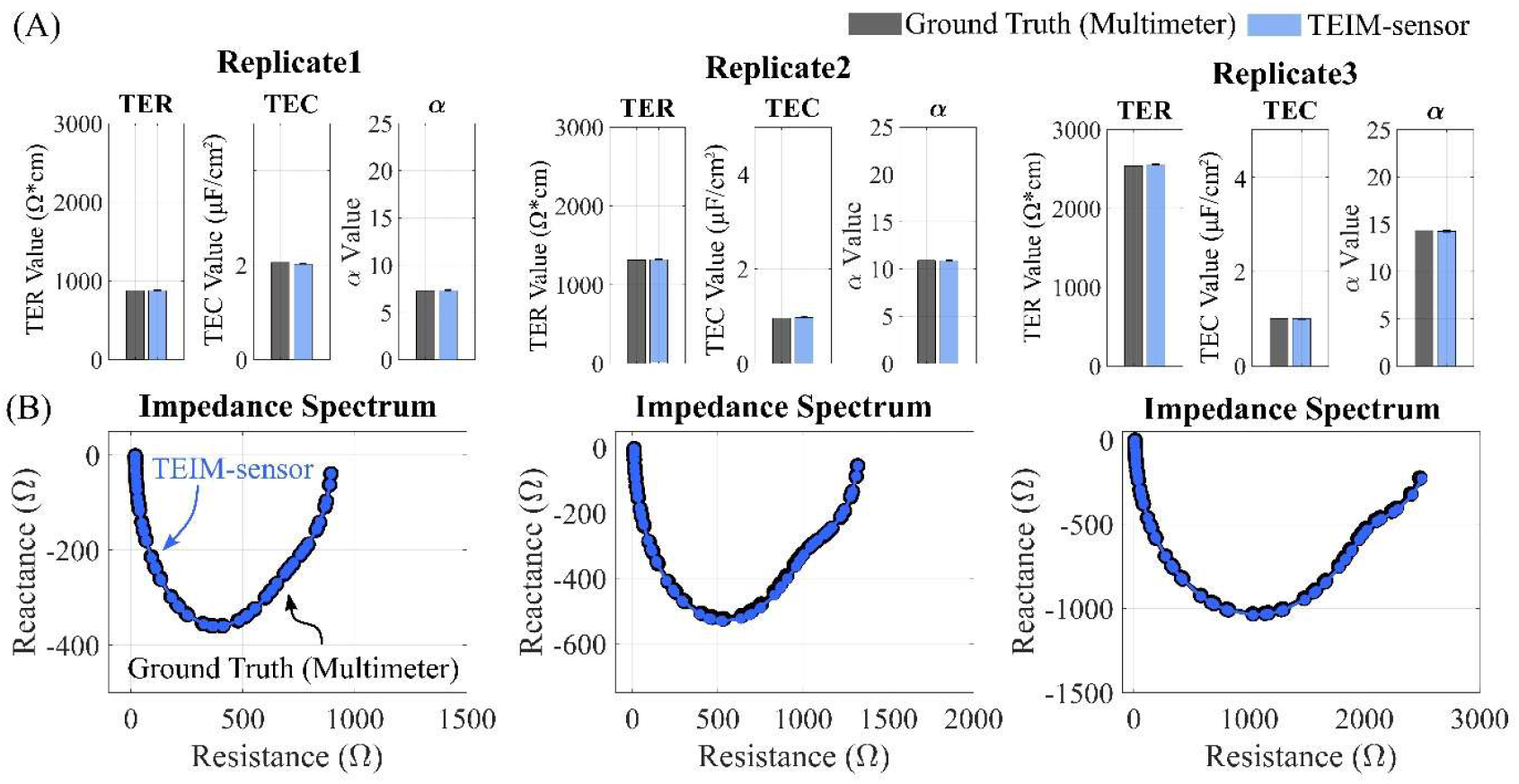
Accuracy and precision of TEIM-sensor on electrical circuit mimics of 16HBE. Data presentation and interpretation are the same as in Figure S2.

